# Modeling individual disability evolution in multiple sclerosis patients based on longitudinal multimodal imaging and clinical data

**DOI:** 10.1101/733295

**Authors:** Ceren Tozlu, Dominique Sappey-Marinier, Gabriel Kocevar, François Cotton, Sandra Vukusic, Françoise Durand-Dubief, Delphine Maucort-Boulch

## Abstract

**Background:** The individual disease evolution of multiple sclerosis (MS) is very different from one patient to another. Therefore, the prediction of long-term disability evolution is difficult based on only clinical information. Magnetic resonance imaging (MRI) provides a very efficient tool to distinguish between healthy and abnormal brain tissue, monitor disease evolution, and help decision-making for personalized treatment of MS patients.

**Objective:** We aim to develop a patient-specific model to predict individual disease evolution in MS, using demographic, clinical, and imaging data that were collected at study onset.

**Methods:** The study included 75 patients tracked over 5 years. The latent class linear mixed model was used to consider individual and unobserved subgroup variability. First, the clinical model was established with demographic and clinical variables to predict clinical disease evolution. Second, the imaging model was built using the multimodal imaging variables. Third, the imaging variables were added one by one, two by two, and all three together to investigate their contribution to the clinical model. The clinical disability is measured with the Expanded Disability Status Scale (EDSS). The performances of the clinical, imaging, and the combined models were compared mainly using the Bayesian Information Criterion (BIC). The mean of the posterior probabilities was also given as the secondary performance evaluation criterion.

**Results:** The clinical model gave higher BIC value than imaging and any combined models. The means of the posterior probabilities given by the three models were over 0.94. The clinical model clustered the patients into two latent classes: stable evolution class (n=6, 88%) and severe evolution class (n=9, 12%).

**Conclusion:** The latent class linear mixed model may provide a well-fitted prediction for the disability evolution in MS patients, thus giving further information for personalized treatment decisions after thorough validation with a larger and independent dataset.

## INTRODUCTION

Multiple sclerosis (MS) is the most frequent disabling neurological disease in young adults. While its etiology remains unknown, MS is a demyelinating, inflammatory, and chronic disease of the central nervous system. The evolution of the disease and the risk of developing permanent disability are very different from one patient to another [Goldenberg, 2012]. Thus, today neurologists’ challenge is to predict the evolution of individual disability using clinical, biological, and imaging data.

MS patients have very different clinical evolution profiles. Further, these profiles may change over time. Clinical impairment is measured with the Expanded Disability Status Scale (EDSS) and its evolution over time is classified currently in four clinical subtypes: clinically isolated syndrome (CIS), relapsing-remittent (RR), primary-progressive (PP), and secondary-progressive (SP). Few patients are early diagnosed as CIS during their first clinical examination. Then, CIS patients may shift to RR that represents 85% of patients. RR patients can shift afterward to SP with or without superimposed relapses [Lawton et al., 2015]. According to Lublin [Lublin, 2014]., 15% of MS patients start with PP that is characterized by continuous worsening of symptoms without relapses since diagnosis period.

In addition to clinical examination, magnetic resonance imaging (MRI) helps in diagnosing MS and monitoring MS evolution. Conventional MRI (such as T1-weighted and T2-weighted imaging) is very sensitive in detecting pathological tissue damage and obtaining valuable predictive information on disease evolution [Filippi, 2001; Fisniku et al., 2008]. Lesion load (LL) and grey matter volume are important markers in evaluating the demyelination level, also axonal and neuronal damage [Peterson et al., 2001; Minneboo et al., 2009]. However, advanced MRI techniques, such as Diffusion Tensor Imaging (DTI) and MR Spectroscopic Imaging, have a better sensitivity and specificity in detecting white. Matter (WM) microstructural damages than conventional imaging [Ge et al., 2004; Sbardella et al., 2013] because of the complementary information based on diffusion and metabolic alterations [Filippi et al., 2001; Filippi, 2001; Filippi, 2001]. Besides, DTI performs well to distinguish MS subtypes because MS patients from different subtypes show specific diffusivity patterns [Sbardella et al., 2013]. Thus, we propose to use DTI data jointly with conventional imaging. Among DTI measurements, fractional anisotropy (FA) measure was chosen because it is very sensitive in detecting microscopic changes related to inflammation [Hannoun et al., 2012].

MS affects white matter tissue and lesions appear mostly around ventricles in periventricular white matter. The floor of the lateral ventricles forms a region called Corpus Callosum (CC), one of the regions that is frequently affected by MS lesions [Barnard, 1974; Gean-Marton et al., 1991; Ge et al., 2006]. The lesions are found in CC in 93% of patients [Gean-Marton et al., 1991]. Moreover, CC is the largest myelinated bundle of the brain providing the connection between the two brain hemispheres. For this reason, the clinical impact of CC lesions is usually more severe compared to other WM lesions. Several studies demonstrated that callosal changes, measured with DTI, were correlated with cognitive and physical disability [Sigal et al., 2012, Yaldizli et al., 2010; Rimkus et al., 2010; Llufriu et al., 2012]. FA measurements were significantly lower in rostrum, body, and splenium parts of CC in MS patients compared to a control group [Hasan et al., 2005; Rueda et al., 2008; Warlop et al., 2008]. The association between EDSS and the atrophy in CC (as measured by conventional MRI) was found in some studies [Hasan et al., 2005; Rueda et al., 2008; Warlop et al., 2008; Schreiber et al., 2001]. However, no significant correlation was found between the disability and callosal atrophy that is measured with the conventional imaging in RR patients [Barkhof et al., 1998]. This contrast can be the result of an insufficiency of conventional imaging of CC; thus, the measurements of CC measured with advanced imaging may be more efficient in MS patients. For this reason, the FA measured in CC was used in the present study.

Several studies have used logistic regression to predict the presence or absence of progression (at least one-point increase in EDSS) or ordinal logistic regression to predict the EDSS changes in ordinal categories [Sastre-Garriga et al., 2005; Minneboo et al., 2008, Khaleeli et al., 2008]. Also, linear regression method was performed to investigate the predictors of the clinical disability using clinical and imaging data. [Minneboo et al., 2009; Furby et al., 2010; Bodini et al., 2011; Enzinger et al., 2011; Popescu et al., 2013]. In addition to these methods, the multilevel approach is performed to consider individual alteration during disability progression to consider heterogeneity among individuals [Di Serio et al. 2009; Lawton et al., 2015]. However, the individual disability evolution in MS does not show a single mean evolution profile; thus, it would be interesting to consider heterogeneity that originated from different mean evolution profiles.

The main objective of the present study is to develop a generalizable predictive model of disability evolution in MS patients, considering unobserved subgroups (different mean-evolution profiles). Therefore, the latent class linear mixed model was used to predict EDSS trajectories over 5 years, using clinical, biological and imaging data collected at the study onset.

## Materials and Methods

### Patients

Eighty patients fulfilling the Mac Donald criteria were included in a standardized clinical and MRI protocol within the frame of the AMSEP project at Lyon Neurological Hospital. This population was divided into 4 groups depending on the MS clinical form: CIS (n=12), RR (n=27), SP (n=16), and PP (n=25). The patients with known ages and disease duration were given after a standardized clinical and MRI examinations every six months during the first three years than at one-year intervals during two years. The clinical examination included EDSS and the Multiple Sclerosis Functional Composite (MSFC) with its three dimensions (Timed 25 Foot Walk [T25FW], 9-Hole Peg Test [9HPT], and Paced Auditory Serial Addition Test (PASAT)). Five patients were excluded from the AMSEP cohort because one or more MSFC component(s) could not be measured at study onset. This left 75 patients for analysis.

### Image Acquisition and Processing

Patients with MS underwent an MR examination on a 1.5T Siemens Sonata system (Siemens Medical Solution, Erlangen, Germany) using an 8-channel head-coil. The MR protocol consisted of in the acquisition of a sagittal 3D-T1 sequence (1 × 1 × 1 mm^3^, TE/TR = 4/2 000 ms) and an axial 2D-spin-echo DTI sequence (TE/TR = 86/6900 ms; 2 × 24 directions of gradient diffusion; b = 1000 s.mm^−2^, spatial resolution of 2.5 × 2.5 × 2.5 mm^3^ oriented in the AC-PC plane).

The lesion load was measured with FLAIR sequence and white and grey matter volumes (GMVs) were measured with T1 imaging without a gadolinium contrast agent.

### The latent class linear mixed model

In our study, the predictions of the individual disability progression were performed with the latent class linear mixed model (LCMM) [Proust-Lima et al., 2015] (See Supplementary Materials for further detail). A linear mixed model assumes that the population is homogeneous and the random effects are normally distributed. However, LCMM considers that the population is not homogeneous and consists of *g* subgroups (also called latent classes). Each latent class shows its distribution with a class-specific matrix of variance-covariance. Each subject belongs to one latent class that maximizes the posterior probability and the sum of probabilities of being in various classes is equal to 1. The posterior probability and Bayesian Information Criterion (BIC) were used to measure the goodness-of-fit of the model [Proust-Lima et al., 2015]. The higher the mean of the posterior probabilities that is obtained at each latent class, the better the classification is.

#### Application of the latent class linear mixed model

Function “hlme” in “lcmm” package of software R was used to implement the latent class linear mixed model. There are three main arguments of the “hlme” function: (1) the **fixed** argument contains the outcome (dependent variable) and the independent variable(s) which has (have) a common effect on all individuals overall latent classes, (2) the **mixture** argument indicates the variable(s) that have a specific effect on each latent class, and (3) the **random** argument contains the variable(s) that have a specific effect at the individual level. All statistical analyses related to the latent class linear mixed model were performed with “lcmm” package in R software version 3.4.0 (2017-04-21).

The EDSS was used as the outcome in the models. The input variables of the models were the clinical (time, age, disease duration, T25FW, 9HPT) and the imaging information (grey matter volume, lesion load, and fractional anisotropy). The EDSS and time were used as longitudinal data. However, the values at study onset were used for the other demographic, clinical and imaging variables.

Before fitting the model, the latent class mixed model requires setting the variable that determines the latent classes and the time function (linear, quadratic, or square root). Time, age, disease duration, T25FW, 9HPT, grey matter volume, lesion load, and fractional anisotropy were tested separately as the variable that determines the latent classes with the linear, quadratic, and square root of time and considering two latent classes. So, we established 24 models and these models were compared using BIC. The model with the lower BIC value provided the best combination of the variable that determines the latent classes and time function.

All demographic, clinical and imaging variables were used in the fixed argument during the choice of the best combination. Besides, age at study onset was used as the variable that exerts the individual effect (in the random argument) in all models because there was a great inter-patient variability in terms of age at study onset.

First, the clinical model was established with age, disease duration, T25FW, 9HPT at study onset, and time in the fixed argument. Second, the imaging model was established with GMV, LL, and FA in the fixed argument. Third, the imaging variables were added one by one, two by two, or all three together with the clinical variables into the clinical model to obtain the combined models. Finally, the clinical, imaging, and combined models were compared using the BIC criterion, knowing that a lower BIC indicates a better fit of the model. Further, AIC, log-likelihood and the mean of the posterior probabilities were also reported in this study to examine the goodness of fit and allow result comparisons with previous studies.

#### Statistical analyses

Because most data did not follow the normal distribution, medians, 1st, and 3rd quartiles were used to describe the data. Consequently, non-parametric statistical tests were used to examine differences in demographic, clinical, and imaging data among the patients of the four clinical subtypes. The Kruskal-Wallis test was used first to check whether the distributions of variables were significantly different. In case of significant difference, a Mann-Whitney U test was used to compare subtype medians two by two. Statistical significance was considered at p<0.05.

## RESULTS

EDSS at study onset was significantly higher for PP and SP patients compared to CIS and RR patients (p-value <0.05). However, there was no significant difference between the EDSS of PP and SP patients (p-value= 0.61). PP patients were significantly older than other patients at study onset (p-value <0.05). The disease duration was similar for RR and PP patients (p-value= 0.51) and was significantly greater for SP patients (p-value <0.05). T25FW was significantly different among all clinical subtypes, and PP patients had the greatest T25FW values at study onset. There was no significant difference between the 9HPT measurements of CIS and RR patients (p-value = 0.11), nor PP and SP patients (p-value =0.63). Also, the GMV was similar for CIS and RR patients (p-value = 0.42) as well as for PP and SP patients (p-value = 0.21). However, the GMV was significantly greater for CIS and RR patients compared to PP and SP patients (p-value <0.05). There was no significant difference in LL between RR and PP patients (p-value=0.86). The LL was significantly lower for CIS patients and greater for SP patients (p-value <0.05). SP patients had the greatest FA value compared to the other clinical subtypes (p-value <0.05).

**Table 1.** Patient demographics, clinical and imaging characteristic.

| **Variable** | **CIS** | **RR** | **PP** | **SP** |
| --- | --- | --- | --- | --- |
| EDSS | 0.50 [0.00, 1.25] | 2.00 [1.25, 4.00] | 4.00 [4.00, 4.50] | 4.50 [4.00, 5.00] |
| Age | 29.00 [28.00, 35.00] | 26.50 [23.25, 34.00] | 36.00 [32.50, 39.50] | 32.00 [24.00, 34.00] |
| Disease Duration | 0.86 [0.27, 2.27] | 4.69 [2.51, 7.76] | 5.66 [4.67, 7.75] | 11.18 [8.14, 15.35] |
| T25FW | 4.15 [3.95, 4.50] | 5.00 [4.20, 6.37] | 6.50 [5.00, 7.70] | 5.90 [5.22, 7.42] |
| 9HPT | 18.85 [17.55, 19.95] | 20.52 [18.06, 23.12] | 24.55 [21.43, 34.88] | 27.23 [23.60, 32.31] |
| GMV | 858.30 [845.70, 882.20] | 841.10 [820.70, 873.40] | 811.20 [781.90, 822.90] | 822.00 [791.50, 832.40] |
| LL | 6.64 [4.28, 8.43] | 14.36 [6.53, 24.32] | 14.33 [11.00, 27.69] | 36.32 [20.31, 41.51] |
| FA | 0.62 [0.60, 0.64] | 0.60 [0.58, 0.62] | 0.59 [0.57, 0.61] | 0.54 [0.48, 0.59] |

*The values at study onset of the demographic, clinical and imaging variables were given and expressed as Median [1<sup>st</sup> quartile, 3<sup>rd</sup> quartile].*

The model with two latent classes that gave the lowest BIC value had 1) the time as the variable with a specific effect on each latent class (in the mixture argument), and 2) a linear time function (in the fixed argument) gave the lowest BIC value (see Appendix). Therefore, in the following sections, the models that included time in the mixture argument and age in the random argument will be discussed

BIC results suggest that the clinical model offers a slightly better fit than the imaging and combined models. All models had mean posterior probabilities of over 0.9. Most of the combined models had a greater mean posterior probability than the clinical and imaging model. The log-likelihood ranged between −550 and −522. The model including the clinical and all imaging variables gave the best likelihood values. AIC values ranged between 1073 and 1125, while the model including clinical and GMV variables had the lowest AIC value.

**Table 2.**
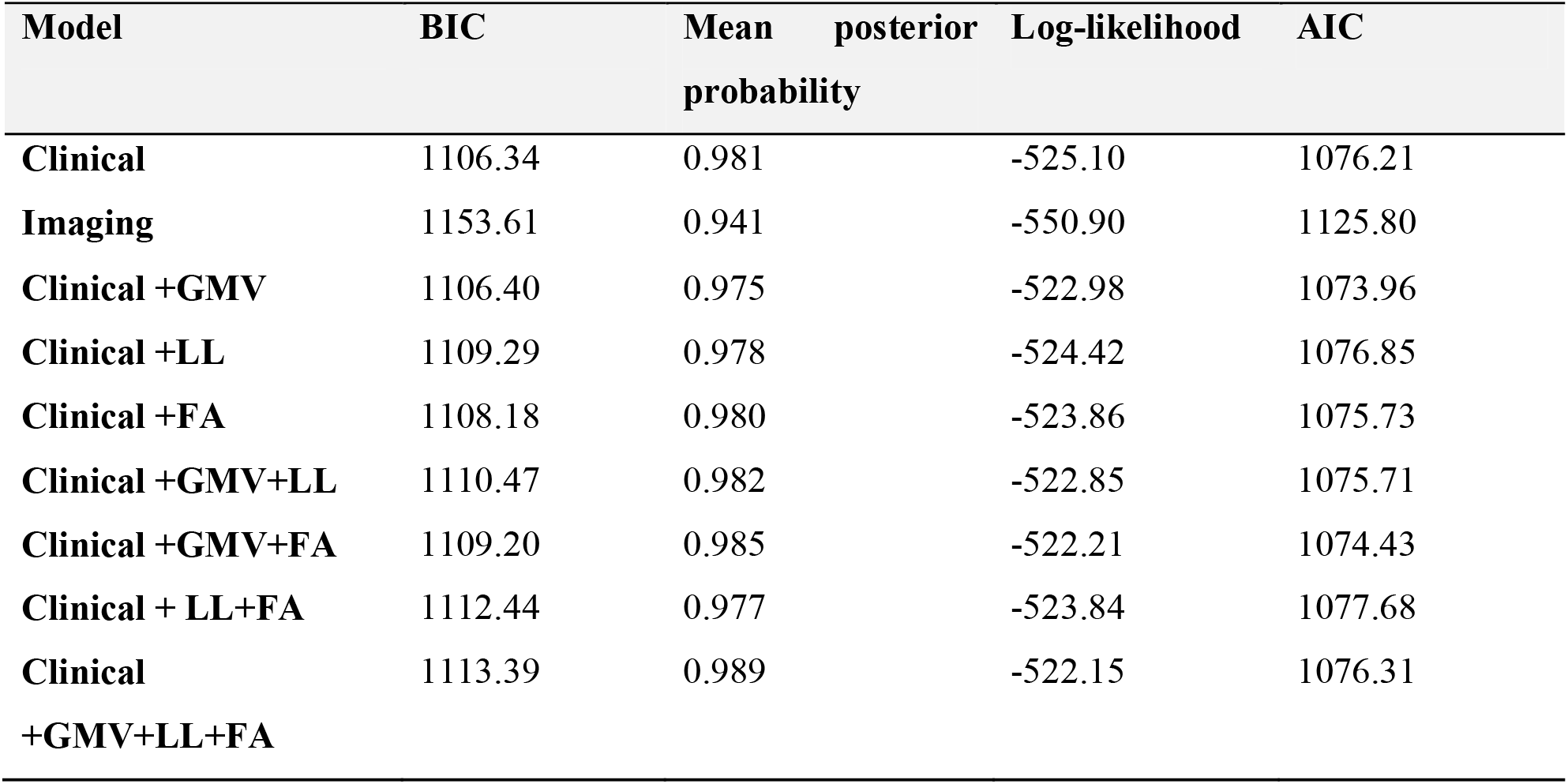
BIC, mean of posterior probabilities, log-likelihood, and AIC results with the clinical, the imaging and the combined models built with different combinations of imaging variables.

| Model | BIC | Mean posterior probability | Log-likelihood | AIC |
| --- | --- | --- | --- | --- |
| **Clinical** | 1106.34 | 0.981 | -525.10 | 1076.21 |
| **Imaging** | 1153.61 | 0.941 | -550.90 | 1125.80 |
| **Clinical +GMV** | 1106.40 | 0.975 | -522.98 | 1073.96 |
| **Clinical +LL** | 1109.29 | 0.978 | -524.42 | 1076.85 |
| **Clinical +FA** | 1108.18 | 0.980 | -523.86 | 1075.73 |
| **Clinical +GMV+LL** | 1110.47 | 0.982 | -522.85 | 1075.71 |
| **Clinical +GMV+FA** | 1109.20 | 0.985 | -522.21 | 1074.43 |
| **Clinical + LL+FA** | 1112.44 | 0.977 | -523.84 | 1077.68 |
| **Clinical +GMV+LL+FA** | 1113.39 | 0.989 | -522.15 | 1076.31 |

The clinical model will be presented hereafter as this model gave the best BIC result. The effect of all clinical was significantly different than zero in the clinical model. The parameter coefficient of the time was given for each latent class and it was significantly different from zero for one of the classes. Specifically, the more time elapsed, the greater the EDSS in Class 1. Also, the parameter coefficients of disease duration, age, T25FW, and 9HPT were positive, and this means that a unit increase of these variables at the study onset had an effect to increase EDSS at the study onset.

**Table 3.**
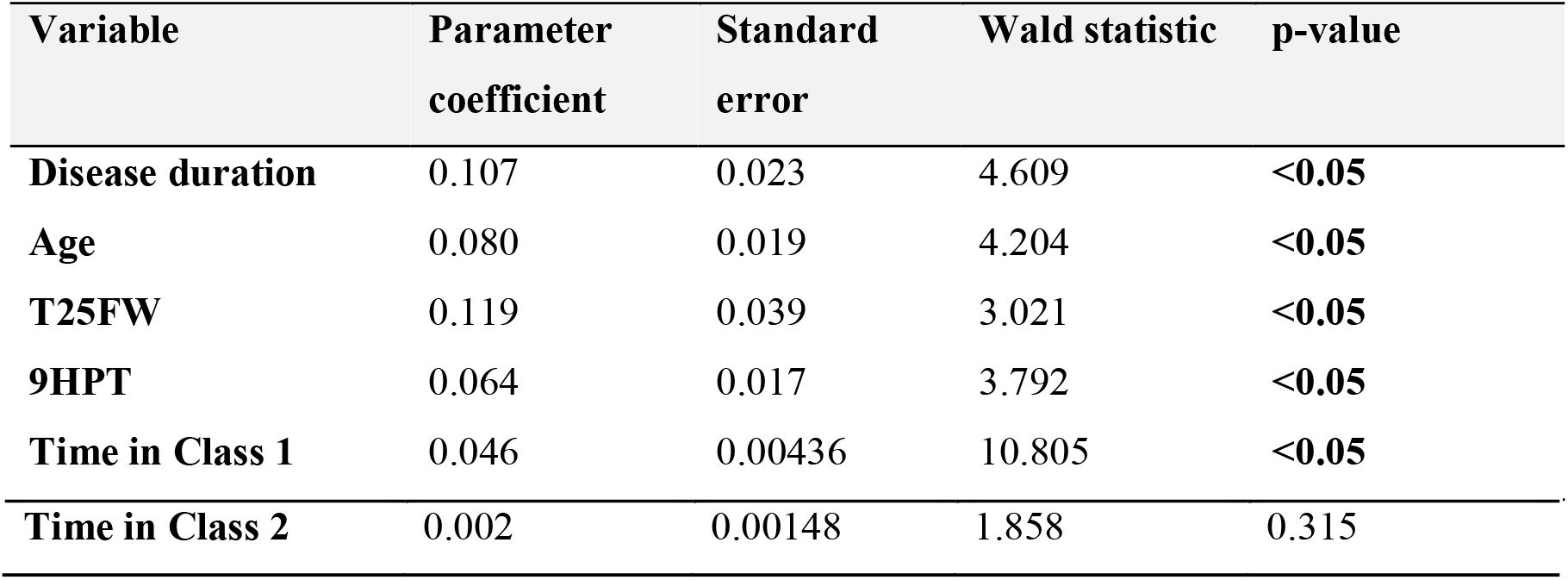
The variables used in the fixed and mixture arguments of the clinical model.

Figure 1 shows the predicted points and observed mean trajectories obtained with the clinical model, and Table 4 gives the classification of the patients in the latent classes. The graph shows that the evolution of EDSS is stable for Class 2. However, Class 1 included all clinical subtypes except CIS and shows a severe evolution of the clinical score. Also, the time was significantly different than zero for this class. Moreover, the predicted points were closer to the observed mean trajectory for Class 2, and Class 1 contained a smaller number of patients compared to Class 2 (12% vs. 88%). The low number of patients in Class 1 might be the cause of the less accurate prediction and the higher difference between the predicted and observed mean trajectories.

**Table 4.** Classification obtained with the clinical model and all MS subtypes.

| Clinical subtype | Class 1 | Class 2 |
| --- | --- | --- |
| CIS | 0 | 12 |
| RR | 2 | 24 |
| PP | 2 | 13 |
| SP | 5 | 17 |

**Figure 1.**
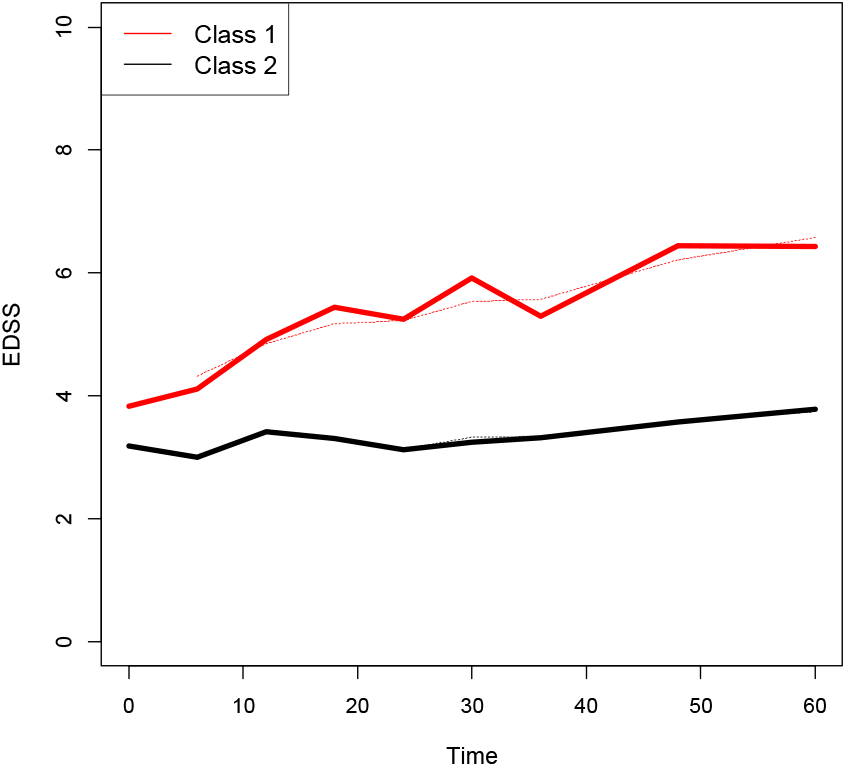
Observed mean trajectories (solid lines) and predicted mean trajectories (dotted lines) of each class according to the clinical model.

The performance of the clinical model was also analyzed based on the prediction at 5 years after the study onset. The predicted and observed EDSS scores were presented in Figure 2. In concordance with the previous results, the patients in Class 1 showed higher predicted and observed EDSS scores compared to the patients in Class 2. The clinical model showed high accuracy with an R^2^ of 0.945 and RMSE of 0.534. The imaging and combined models gave similar results with an R^2^ over 0.9 and RMSE of less than 0.6. The prediction was poor for the patients with lower EDSS scores (i.e. lower than EDSS 4). The patients with lower EDSS scores might be classified in another group that would lead to a better prediction.

**Figure 2.**
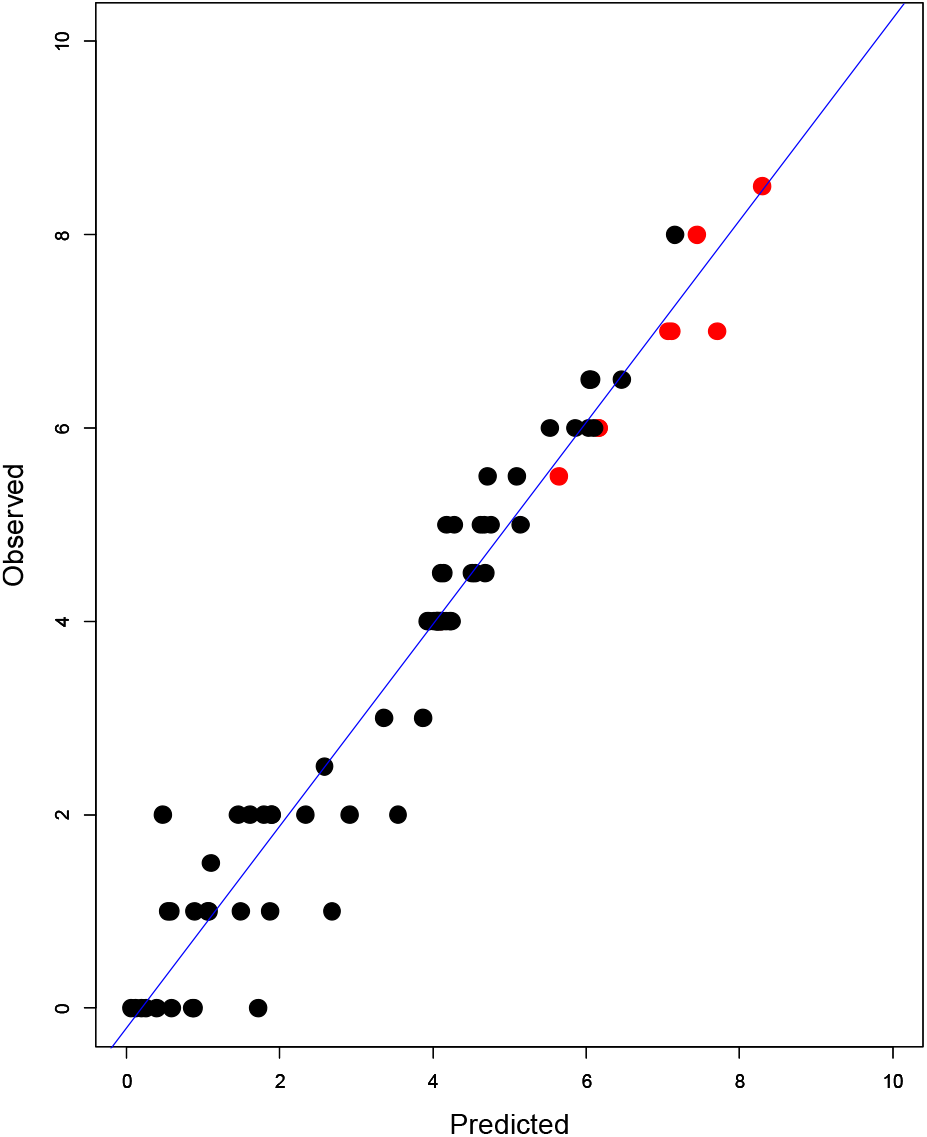
Observed and predicted EDSS at 5 years after the study onset for Class 1 (red) and Class 2 (black) according to the clinical model. The blue line which was obtained with a linear regression model was also given on the figure.

The patients in the two latent classes were compared based on EDSS, age, disease duration, T25FW, 9HPT, GMV, LL, and FA at study onset. Table 5 shows that the LL was significantly greater in the class, in which the patients show an aggressive disability progression (Class 1) (p-value<0.05). However, other demographic, clinical, and imaging variables were not significantly different between the two latent classes (p-value > 0.05 for all variables). Figure 3 demonstrates that the LL in Class 1 showed a normal distribution, whereas the LL was asymptotically distributed in Class 2. Most of the patients in Class 2 had less LL compared to Class 1. Also, the median of the LL in Class 1 was approximately two times higher than the median of LL in Class 2.

**Table 5.** Clinical and imaging variables according to each latent class obtained with the clinical model.

| **Variable** | **Class 1**<br>**N=9** | **Class 2**<br>**N=66** | **p-value** |
| --- | --- | --- | --- |
| **EDSS** | 4.00 [4.00, 4.50] | 4.00 [2.00, 4.38] | 0.31 |
| **Age** | 40.510 [32.470, 44.080] | 39.120 [30.410, 42.850] | 0.64 |
| **Disease duration** | 6.694 [4.991, 10.582] | 5.798 [2.314, 8.813] | 0.34 |
| **9HPT** | 24.85 [20.10, 32.65] | 22.10 [19.35, 26.34] | 0.41 |
| **25FW** | 5.050 [4.100, 7.000] | 5.325 [4.500, 6.588] | 0.60 |
| **GMV** | 831.1 [799.8, 850.7] | 825.4 [804.4, 859.5] | 0.71 |
| **LL** | 26.446 [15.052, 35.412] | 14.296 [6.412, 32.469] | <0.05 |
| **FA** | 0.578 [0.556, 0.621] | 0.599 [0.5796, 0.623] | 0.48 |

*The values are expressed as Median [1st, 3rd quartile]. The p-value is the result of Kruskal-Wallis test which was performed to compare the distribution of the variable values in two classes.*

**Figure 3.**
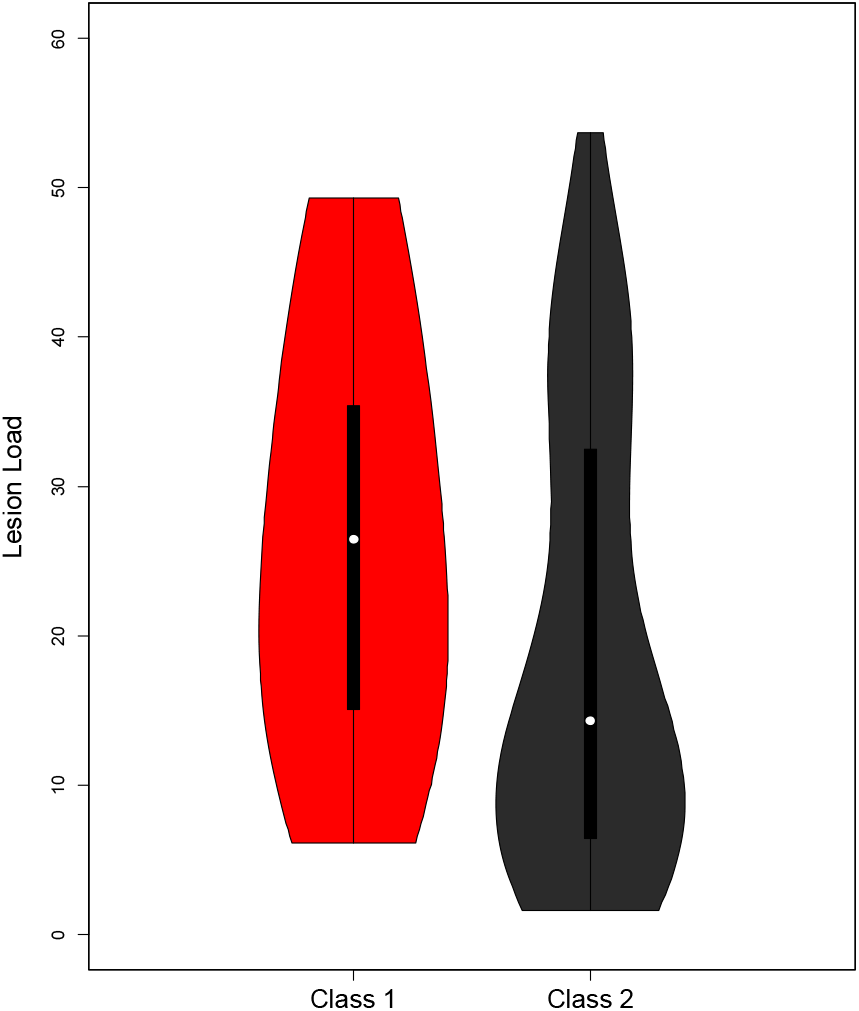
The violin plot of LL which was observed in Class 1(red) and Class 2 (black). The classes were established by the clinical model. The violin plot shows the density, median, 1^st^ and 3^rd^ quartile of LL.

### Imaging and combined models

Table 6 shows that the effect of the GMV was significantly different from zero (p-value <0.05) in the imaging model but not significant (p-value=0.062) in the combined model. Time in Class 1, as well as the demographic and clinical variables, were still significant in the combined model (p-value <0.05). However, the imaging variables did not have any significant effect, and the parameter coefficients of these parameters were not significantly different than zero (p-value_GMV_=0.062, p-value_LL_=0.723, and p-value_FA_=0.231). The EDSS evolution trajectories and the classification of the patients that were found by imaging and combined models were similar to the results of the clinical model (See Supplementary Material). There was only one more SP patient classified in Class 2 (with stable evolution) instead of Class 1 (with severe evolution) in the combined model results.

**Table 6.** The results of the imaging and combined models.

| **Imaging Model** |  |  |  |  |
| --- | --- | --- | --- | --- |
| **Variable** | **Parameter coefficient** | **Standard error** | **Wald statistic** | **p-value** |
| GMV | -0.020 | 0.004 | -4.648 | **<0.05** |
| LL | -0.008 | 0.016 | -0.529 | 0.596 |
| FA | -0.402 | 0.288 | -1.394 | 0.163 |
| Time in Class 1 | 0.047 | 0.004 | 11.213 | **<0.05** |
| Time in Class 2 | 0.002 | 0.002 | 1.027 | 0.304 |

| **Combined model** |  |  |  |  |
| --- | --- | --- | --- | --- |
| Disease duration | 0.103 | 0.023 | 4.491 | **<0.05** |
| Age | 0.059 | 0.022 | 2.660 | **<0.05** |
| T25FW | 0.124 | 0.036 | 3.396 | **<0.05** |
| 9HPT | 0.042 | 0.019 | 2.220 | **<0.05** |
| GMV | -0.008 | 0.005 | -1.860 | 0.062 |
| LL | -0.004 | 0.011 | -0.354 | 0.723 |
| FA | -0.258 | 0.216 | -1.196 | 0.231 |
| Time in Class 1 | 0.047 | 0.004 | 11.553 | **<0.05** |
| Time in Class 2 | 0.002 | 0.002 | 1.173 | 0.240 |

## DISCUSSION

The latent class linear mixed model was used to model the evolution of disability in patients with multiple sclerosis, and thus, to predict individual disability evolution as measured by the EDSS. To our knowledge, this is the first study that predicts long-term disability evolution considering unobserved classes of patients with multiple sclerosis from all clinical subtypes. First, the clinical and the imaging models were separately built with clinical and imaging variables, respectively. Then, multimodal imaging variables were added to the clinical model to obtain combined models. As per the BIC criterion, the clinical model had higher predictive accuracy in comparison with the imaging or the combined models.

In the clinical and combined models, all clinical variables had a significant predictive effect on disease evolution. Previous studies also showed significant effects of age and disease duration in predictive modeling [Confavreux et al., 2003; Scalfari et al., 2011]. Besides, 9HPT and T25FW were considered two of the best measures across a wide range of indicators of MS disability [Kieseier & Pozzilli, 2011; Kraft et al., 2014]. However, no previous study had checked whether the 9HPT had a significant effect on disability evolution. On the other hand, an early change in T25FW was significant to the long-term EDSS evolution in progressive patients (PP and SP) [Bosma et al., 2012]. Our results confirmed that the effects of T25FW and 9HPT were significant in all clinical subtypes.

Previous studies on predictive modeling of disability evolution in patients with multiple sclerosis used logistic regression models, Kaplan-Meier analyses, Markov models, multilevel modeling, and, mostly, linear mixed models [Popescu et al., 2013; Minneboo et al., 2009; Bodini et al., 2011; Minneboo et al., 2007; Palace et al., 2014; Lawton et al., 2015; Confavreux et al., 2003; Confavreux et al., 2000]. A recent study that used latent class linear mixed models in PP patients to identify unobserved classes [Signori et al., 2017] selected the best model with the BIC criterion and identified three subgroups of PP patients. However, our study remains the first one to use latent class mixed models, consider unobserved classes, and include all clinical subtypes of MS.

In our study, patients with multiple sclerosis were classified into two latent classes (stable and severe evolution over time). Most of the patients (88%) showed no progression of disability over five years after the study onset. This observation was not surprising as all patients were taking medication, which was effective in most cases. Further, MS is a long-lasting disease, and the five-year study period might have been insufficient to show disability worsening. However, 12% of the patients were assigned to the class with severe disability progression. Assigning patients to either of these two latent classes (stable or severe) may help treatment decisions. For example, patients with a probable severe evolution would benefit from second-line therapies.

EDSS 4 is known to be the threshold of limited walking disability, even though a patient may be able to walk more than 500 m. The predicted EDSS values were around EDSS 4 at the study onset in the three latent classes, and then each class evolved differently. Thus, our predictions were able to distinguish the patients who might have a severe, stable, or moderate evolution after reaching the threshold of limited walking disability.

GMV had a significant role in the imaging model in our study. Indeed, previous studies have shown that GMV may constitute a good marker of the risk of increased disability in MS patients [Rovaris et al., 2006; Fisniku et al., 2008, Roosendaal et al., 2011; Sepulcre et al., 2006]. However, the effects of the multimodal imaging variables were not statistically significant in the combined models. Enzinger et al. have performed multivariate analysis and found that no MRI variable had a significant effect on disability evolution [Enzinger et al., 2011]. In contrast, Popescu et al. showed that the LL measured in grey and white matter was a strong predictor of the 10-year EDSS in MS [Popescu et al., 2013]. However, the white matter LL measured in our study did not play a significant role, neither in the imaging nor in the combined models.

One limitation of the present work was the use of the LL measured in the whole WM rather than locally. The location and size of the lesion are very important markers of clinical disability. A future modeling study on the evolution of clinical disability may consider the focal LL in WM and GM.

To conclude, the latent class linear mixed model allowed building a well-fitted predictive model for disability evolution in patients with MS, and this model showed highly accurate results. The model developed here is highly promising in predicting individual long-term disability evolution in all clinical subtypes of MS.

## Acknowledgements

The work of the first author was supported by Open Health Institute.

The authors thank Jean Iwaz (Hospices Civils de Lyon) and Nathan Lindberg (Cornell University) for the revision of the final drafts of the manuscript.

## Author Contribution statement

CT carried out the statistical analyses and drafted the article.

DSM acquired, organized, checked, and managed the imaging data, and commented and reviewed the article.

GK organized and managed the imaging data, and commented and reviewed the article.

FDD and SV carried out clinical data collection and commented and reviewed the article.

DMB organised the study and commented and reviewed the article.

