## Supplementary document for "Modeling individual disability evolution in multiple sclerosis patients based on longitudinal multimodal imaging and clinical data"

**SUPPLEMENTARY MATERIAL**

***Supplementary Table 1 -*** *Bayesian Information Criterion (BIC) and Akaike Information Criterion (AIC) obtained with 24 models fitted with 8 different variables (time, age, disease duration, 9HPT, and T25FW) in mixture argument, and 3 types of time function (linear, square root, or polynomial) considering 2 latent classes.*

| Variable in the mixture argument | Time function | BIC | AIC |
| --- | --- | --- | --- |
| Time | Linear | 1113.394 | 1076.314 |
|  | Square Root | 1117.601 | 1078.204 |
|  | Polynomial | 1217.116 | 1177.719 |
| Age | Linear | 1196.412 | 1159.332 |
|  | Square Root | 1209.360 | 1169.963 |
|  | Polynomial | 1200.156 | 1160.759 |
| Disease Duration | Linear | 1213.291 | 1176.212 |
|  | Square Root | 1201.165 | 1161.767 |
|  | Polynomial | 1215.503 | 1176.106 |
| 9HPT | Linear | 1207.134 | 1170.054 |
|  | Square Root | 1211.055 | 1171.658 |
|  | Polynomial | 1210.959 | 1171.562 |
| T25FW | Linear | 1213.291 | 1176.212 |
|  | Square Root | 1205.761 | 1166.364 |
|  | Polynomial | 1208.620 | 1169.223 |
| GMV | Linear | 1213.291 | 1176.212 |
|  | Square Root | 1201.102 | 1161.704 |
|  | Polynomial | 1200.969 | 1161.571 |
| LL | Linear | 1195.025 | 1157.945 |
|  | Square Root | 1212.858 | 1173.460 |
|  | Polynomial | 1198.706 | 1159.309 |
| FA | Linear | 1192.264 | 1155.184 |
|  | Square Root | 1211.315 | 1171.918 |
|  | Polynomial | 1195.941 | 1156.544 |

Imaging Model

The predicted and observed mean trajectories obtained with the imaging model are presented on Figure 2. The predicted trajectories were similar to the trajectories obtained with the clinical model. Class 2 trajectory included the majority of patients (88%) and showed a stable evolution. Class 1 included two RR, two PP, and four SP patients. The predicted mean trajectories of these patients showed severe evolutions.

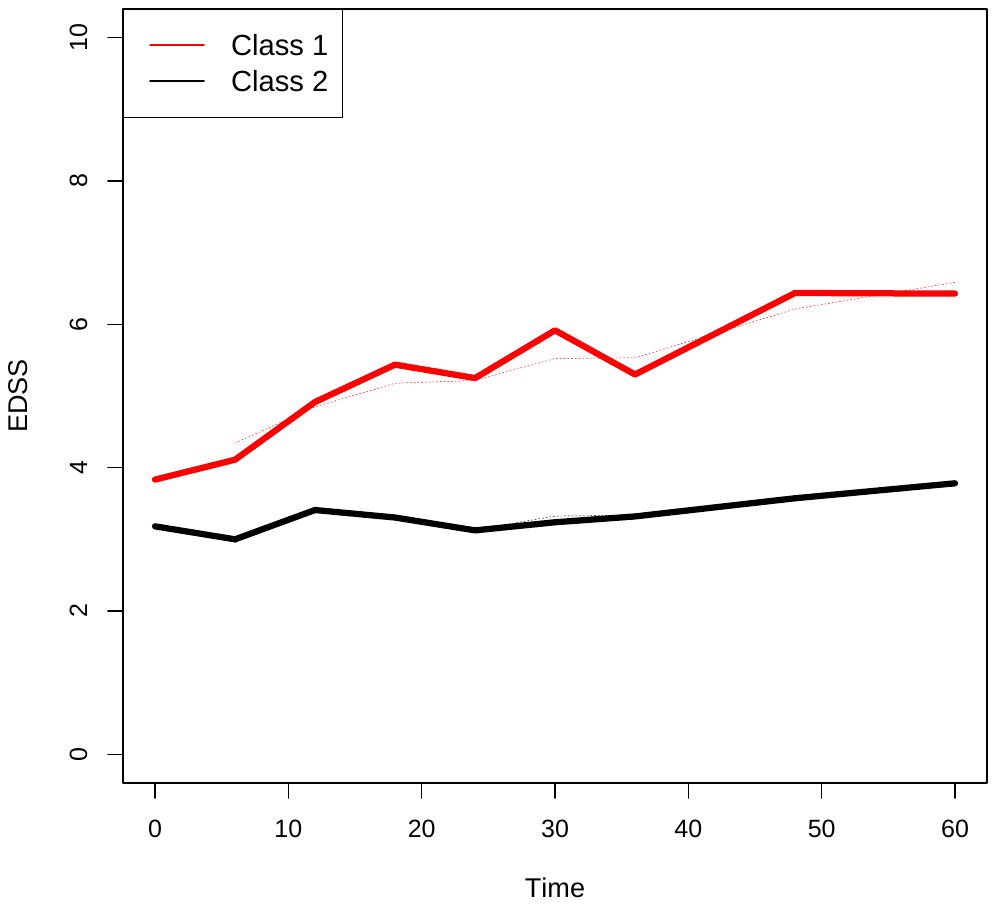

***Supplementary Figure 1 -*** *Observed mean trajectories (solid lines) and predicted mean trajectories (dotted lines) of each class according to the imaging model.*

***Supplementary Table 2 -*** *Classification results of the imaging model.*

| **Clinical subtype** | **Class 1** | **Class 2** |
| --- | --- | --- |
| **CIS** | 0 | 12 |
| **RR** | 2 | 24 |
| **PP** | 2 | 13 |
| **SP** | 5 | 17 |

Combined Model

Figure 3 shows the predicted and observed mean trajectories obtained with the combined model built with GMV, LL, and FA together. The observed mean trajectories and the predicted points were almost similar to the result obtained with the clinical data. Moreover, the classification performed with the combined model (see Table 8) did not significantly change the classification obtained with the clinical model.

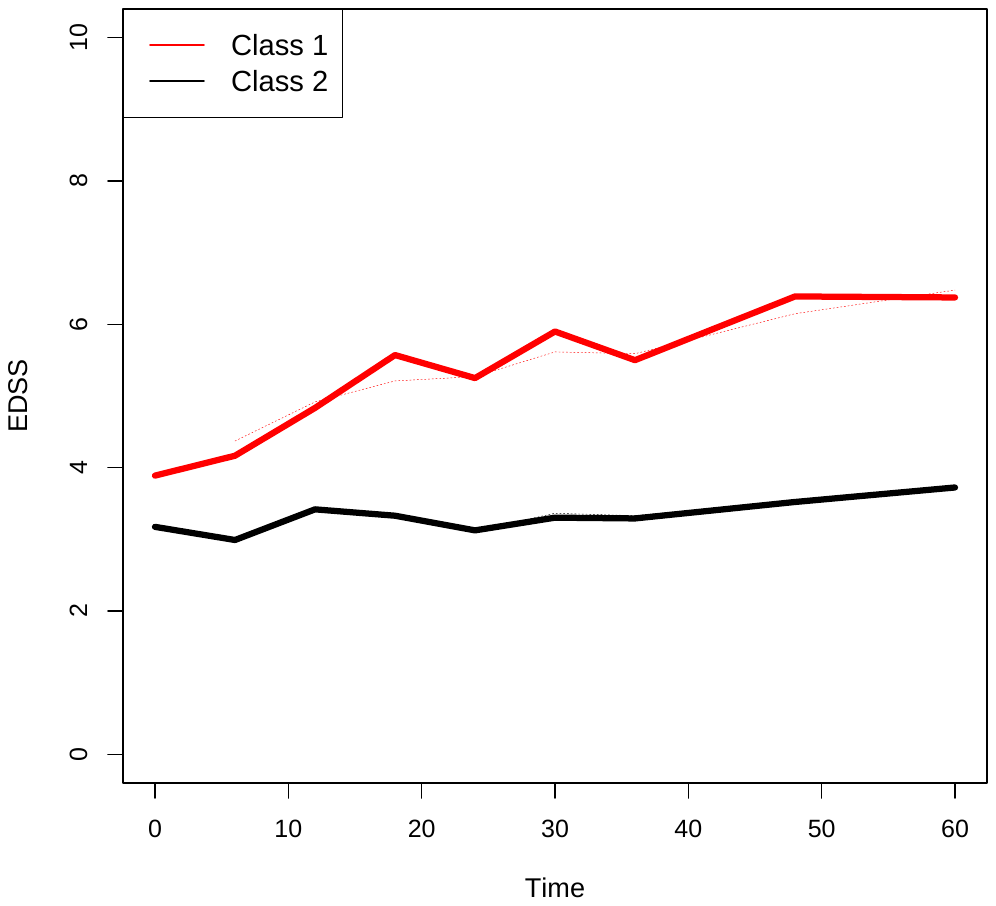

***Supplementary Figure 2 -*** *Observed mean trajectories (solid lines) and predicted mean trajectories (dotted lines) of each class according to the combined model.*

***Supplementary Table 3 -*** *Classification results of the combined model (demographic, clinical*

*variables with GMV, LL and FA).*

| **Clinical subtype** | **Class 1** | **Class 2** |
| --- | --- | --- |
| **CIS** | 0 | 12 |
| **RR** | 2 | 24 |
| **PP** | 2 | 13 |
| **SP** | 4 | 18 |
